# Efficient transposon mutagenesis mediated by an IPTG-controlled conditional suicide plasmid

**DOI:** 10.1101/419473

**Authors:** Santa S. Naorem, Jin Han, Stephanie Y. Zhang, Junyi Zhang, Lindsey B. Graham, Angelou Song, Cameron V. Smith, Fariha Rashid, Huatao Guo

## Abstract

**Background:** Transposon mutagenesis is highly valuable for bacterial genetic and genomic studies. The transposons are usually delivered into host cells through conjugation or electroporation of a suicide plasmid. However, many bacterial species cannot be efficiently conjugated or transformed for transposon saturation mutagenesis. For this reason, temperature-sensitive (*ts*) plasmids have also been developed for transposon mutagenesis, but prolonged incubation at high temperatures to induce *ts* plasmid loss can be harmful to the hosts and lead to enrichment of mutants with adaptive genetic changes. In addition, the *ts* phenotype of a plasmid is often strain- or species-specific, as it may become non-*ts* or suicidal in different bacterial species.

**Results:** We have engineered several conditional suicide plasmids that have a broad host range and whose loss is IPTG-controlled. One construct, which has the highest stability in the absence of IPTG induction, was then used as a curable vector to deliver hyperactive miniTn5 transposons for insertional mutagenesis. Our analyses show that these new tools can be used for efficient and regulatable transposon mutagenesis in *Escherichia coli, Acinetobacter baylyi* and *Pseudomonas aeruginosa*. In *P. aeruginosa* PAO1, we have used this method to generate a Tn5 insertion library with an estimated diversity of ~10^8^, which is ~2 logs larger than the best transposon insertional library of PAO1 and related *Pseudomonas* strains previously reported.

**Conclusion:** We have developed a number of IPTG-controlled conditional suicide plasmids. By exploiting one of them for transposon delivery, a highly efficient and broadly useful mutagenesis system has been developed. As the assay condition is mild, we believe that our methodology will have broad applications in microbiology research.

## Background

Transposon mutagenesis is a powerful technique for bacterial genetic and genomic studies. One of the most widely used transposons is derived from Tn5. The Tn5 transposon contains two IS50 elements as inverted terminal repeats (Additional file 1: Figure S1) [1, 2]. Both IS50 and Tn5 can be mobilized by their encoded transposase (Tnp) protein, which recognizes two 19 base pair (bp) sequences at their ends, namely outside end (OE) and inside end (IE), for transposition [2]. OE and IE differ by 7 bp (Additional file 1: Figure S1). As Tn5 insertion is almost completely random, it can insert into any gene in a bacterium. The native Tn5/IS50 is not very active, thus avoiding overt deleterious effect on their hosts, but hyperactive mutants have been engineered as genetic manipulation tools [2, 3]. The most active one contains a mosaic sequence of OE and IE (mosaic end; ME) at the transposon termini and an engineered *tnp* gene encoding a highly active transposase enzyme (Tnp^H^), which together increase Tn5 transposition by more than 1000-fold.

Transposons for insertional mutagenesis are usually delivered into bacteria through conjugation of a suicide plasmid [4-6]. Insertion mutants are then selected as the transposons are tagged with an antibiotic-resistance gene. The success of a transposon mutagenesis assay, especially a saturation mutagenesis assay, requires generation of an insertion library with high diversity, which requires efficient plasmid conjugation and transposon transposition. However, conjugation is inefficient in many bacterial species. Occasionally, electroporation has also been used to deliver transposon-containing suicide plasmids for mutagenesis, but low library diversities were often achieved using such approaches [7-9]. To perform efficient transposon mutagenesis in these organisms, temperature-sensitive (*ts*) plasmids are sometimes used for transposon delivery [10-16]. However, many organisms do not have a *ts* and easily manipulatable plasmid, and sometimes a *ts* plasmid in one organism is either non-*ts* or suicidal in a different organism [10, 14, 17]. In addition, a high temperature is often required to cure the *ts* plasmids after mutagenesis, which can be inhibitory to cell growth and may result in selection of mutants with adaptive genetic changes [10, 11, 14].

In this study, we have developed an efficient and regulatable transposon mutagenesis tool that exploits an IPTG-controlled conditional suicide plasmid. It contains an RSF1010 replicon, an IncQ-type replication origin that allows plasmid replication in most Gram-negative bacteria, as well as a few Gram-positive bacteria [18]. It is relatively small, so it can be easily modified. To control plasmid replication by IPTG, a second copy of the plasmid-encoded *repF* repressor gene is cloned downstream of the *Escherichia coli tac* promoter. For efficient and regulatable transposon mutagenesis, we used miniTn5 (mTn5) transposons and cloned the hyperactive transposase gene downstream of a *lac* promoter. We show that the resulting constructs can be used for efficient insertional mutagenesis in three different bacterial species. In *Pseudomonas aeruginosa* PAO1, we show that our system is able to generate a Tn5 insertion library that is almost 2 logs larger than the best library of PAO1 and related *Pseudomonas* strains previously reported, demonstrating that we have developed a powerful mutagenesis tool that is highly useful for microbiology studies.

## Results

### Construction of IPTG-controlled suicide plasmids

To develop a method for efficient transposon mutagenesis in bacterial species that are difficult to transform and conjugate, we created multiple IPTG-controlled suicide plasmids that have a broad host range (Fig. 1a). The plasmids were derived from pMMB208, which is a conjugatable plasmid containing an RSF1010 *oriV* (an IncQ-type origin of replication) that can replicate in most Gram-negative bacteria and a few Gram-positive bacteria [18]. Plasmid replication requires three proteins, RepA, MobA/RepB and RepC, which are a helicase, a primase and an *oriV*-binding protein, respectively. *repF* encodes a small repressor protein that binds the P4 promoter and controls the *repF*-*repA*-*repC* operon through feedback inhibition [19, 20]. pMMB208 also contains a *tac* promoter (P*tac*), a *lacI*^*Q*^ gene and a chloramphenicol resistance marker (*Cam*^*R*^). To create a conditional suicide plasmid (pMMB-*repF*), a second copy of the *repF* gene was inserted downstream of P*tac*. Upon IPTG induction, efficient plasmid loss from transformed *E. coli* DH10B cells was observed (99.97%; Fig. 1b). As an alternative strategy, we inserted two *repA* helicase dominant negative mutants, K42A and D139A, downstream of P*tac* [21]. Similarly, IPTG was able to induce efficient plasmid loss from the transformed DH10B cells. In fact, plasmid retention rates of the dominant negative mutants (K42A, 4.9 x 10^−7^; D139A, 1.5 x 10^−5^) were much lower than that of pMMB-*repF* (3.5 x 10^−4^) (Fig. 1b). However, the two *repA* dominant negative mutants showed significantly lower plasmid stability in the absence of IPTG induction (Fig. 1b), suggesting that plasmid replication is strongly inhibited by leaky expression of the dominant negative mutants, or by spontaneous recombination of the wild-type and the dominant negative *repA* genes (~861 bp direct repeats). Consistent with that, there were ~20-30-fold less plasmid isolated from the same amount of cells for the two mutant constructs (Fig. 1c). Therefore, we decided to choose pMMB-*repF* for further experiments. To kill the cells that still retain the plasmid after IPTG induction, we inserted a *sacB* counter selection marker into the vector [22], resulting in pMMB-*repF*/*sacB*. Indeed, insertion of *sacB* allows efficient killing of plasmid-containing cells by sucrose (data not shown; also see below).

**Fig. 1.**
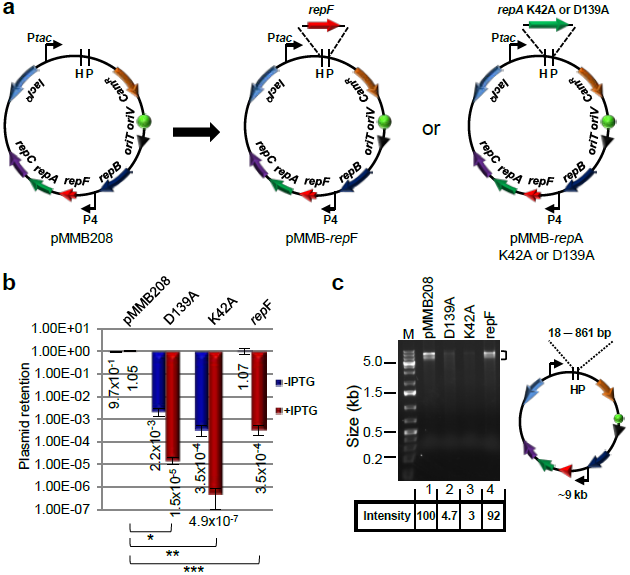
IPTG-controlled conditional suicide plasmids. (**a**) Plasmid pMMB208 and its conditional-suicide derivatives. pMMB208 contains an RSF1010 *oriV* for replication and an *oriT* for conjugation. Genes *repA, mobA/repB* and *repC* encode proteins required for plasmid replication, and *repF* encodes a transcription repressor that binds promoter P4. pMMB208 also has *Cam*^*R*^ and *lacI*^*Q*^ genes and a P*tac* promoter. Plasmid pMMB-*repF* is a derivative of pMMB208 that has a second copy of the *repF* gene inserted downstream of P*tac*. Plasmids pMMB-*repA*K42A and pMMB-*repA*D139A have a dominant-negative *repA* mutant gene, either K42A or D139A, inserted downstream of P*tac*. (**b**) Amount of *E. coli* DH10B cells retaining the indicated plasmids after 24 h growth in the absence of antibiotics, either with or without IPTG induction. Results were average of three independent experiments, and bars represent mean ± SD (standard deviation). ^*^p < 0.0001, ^**^p < 0.0001, and ^***^p < 0.0001 by unpaired Student’s t-test for IPTG induced cultures. (**c**) pMMB208 and its derivatives are digested with *Hin*dIII (H) and *Pst*I (P). Comparing to pMMB208 and pMMB-*repF*, the *repA* K42A and D139A mutants showed reduced yields in plasmid minipreps (no IPTG induction; 3.0% and 4.7% of that of pMMB208, respectively). *Hin*dIII and *Pst*I digestion generates two fragments for each plasmid. The ~9 kb fragment is seen on the gel, while the shorter ones, ranging from 18 bp for pMMB208 to 861 bp for the *repA* mutants, are not visible. Another large band (~9 kb) is also seen in restriction digestion of pMMB208 and its derivatives, even after complete digestion, and the cause is unknown.

### IPTG-controlled mutagenesis of *E. coli* by a highly-active mTn5 transposon

A *Kan*^*R*^-tagged mTn5 was then inserted in pMMB-*repF*/*sacB* for transposon mutagenesis (Fig. 2a) [4]. The mTn5 contains an OE and an IE at the termini. In addition, it contains an uncoupled, *lac* promoter (P*lac*)-controlled *tnp*^*H*^ gene encoding the hyperactive transposase (Tnp^H^) [3], thus allowing inducible expression of Tnp^H^. *E. coli* cells transformed with this plasmid, pSNC-mTn5, were cultured in LB media with and without IPTG induction for 24 h. Cells were then analyzed for efficiencies of plasmid loss, sucrose counter selection and transposon insertion (See Materials and Methods). The plasmid is stable without IPTG induction, as ~91.4% of cells retained the plasmid (Cam^R^) after 24 h culture in the absence of antibiotics (Fig. 2b). In contrast, ~2.6 x 10^−3^ of the cells retained the plasmid post IPTG induction, suggesting that overexpression of the RepF repressor caused efficient plasmid loss. Sucrose counter selection further reduced plasmid-bearing cells (~1.6 x 10^−6^ are Cam^R^; ~1600-fold reduction). In comparison, the percentage of Suc^R^Kan^R^ cells after IPTG induction was found to be ~2.3 x 10^−4^ (Tn5-containing), significantly higher than that of Suc^R^Cam^R^ cells (~1.6 x 10^−6^, plasmid-containing), suggesting that Tn5 transposition had occurred efficiently (Suc^R^Kan^R^Cam^S^: ~2.3 x 10^−4^). Colony restreaking showed that 150/150 Suc^R^Kan^R^ colonies were Kan^R^Cam^S^ (Fig. 2c). Colony PCRs, which used two sets of primers (P1+P2 for detection of *Kan*^*R*^, or mTn5, and P3+P4 for detection of *tac*-*repF*, or plasmid), confirmed plasmid loss in 10 out of 10 colonies (10/10) (Fig. 2d). Sequence analysis showed that all 13 Suc^R^Kan^R^ colonies analyzed had different Tn5 insertion sites (Fig. 3a).

**Fig. 2.**
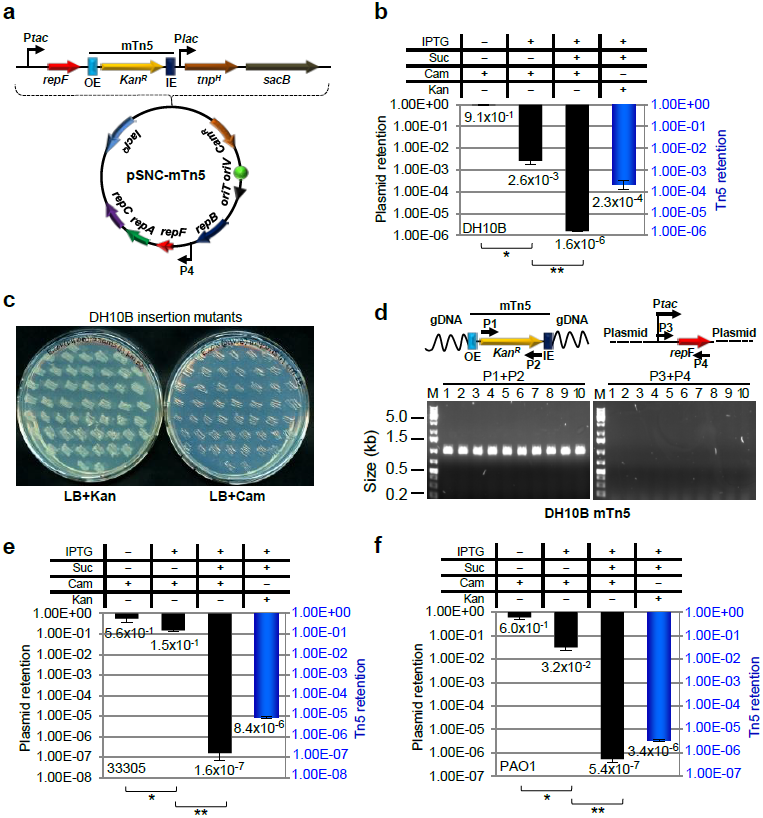
mTn5 transposon mutagenesis using an IPTG-controlled conditional suicide plasmid. (**a**) Diagram of plasmid pSNC-mTn5. pSNC-mTn5 is a derivative of *pMMB-repF* that contains a *Kan*^*R*^-tagged mTn5, a *lac* promoter-controlled hyperactive transposase gene (*tnp*^*H*^), and a *sacB* counter selection marker (with its own promoter). OE and IE are outside and inside ends of the mTn5. (**b**) Plasmid and transposon retention frequencies in *E. coli* DH10B. A “+” symbol for IPTG indicates that the inducer was added to the liquid culture, and a “+” symbol for Suc, Cam, and Kan indicates that the chemicals were added to the plates. Black columns represent plasmid retention frequencies, and the blue column represents Tn5 retention frequency. Results were average of three independent experiments, and bars represent mean ± SD (^*^p < 0.0001 and ^**^p = 0.0054 by unpaired t-test). (see Materials and Methods for details) (**c**) Colony restreaking. 150/150 Suc^R^Kan^R^ colonies of DH10B were found to be Kan^R^Cam^S^ and 50 are shown here. (**d**) Colony PCR of 10 restreaked clones in (**c**). Primer sets P1&P2 and P3&P4 detect *Kan*^*R*^ and *repF*, respectively. All were mTn5-positive and plasmid-negative. Primers P3 and P4 are a functional pair for PCR-amplification of the plasmid sequence (data not shown). (**e**) Plasmid and transposon retention frequencies in *A. baylyi.* Results were average of three independent experiments, and bars represent mean ± SD (^*^p = 0.024 and ^**^p < 0.0001 by unpaired t-test). (**f**) Plasmid and transposon retention frequencies in *P. aeruginosa*. Results were average of three independent experiments, and bars represent mean ± SD (^*^p = 0.0013 and ^**^p = 0.0038 by unpaired t-test). Colony restreaking and PCR analysis are shown in Additional file 1: Figure S2.

**Fig. 3.**
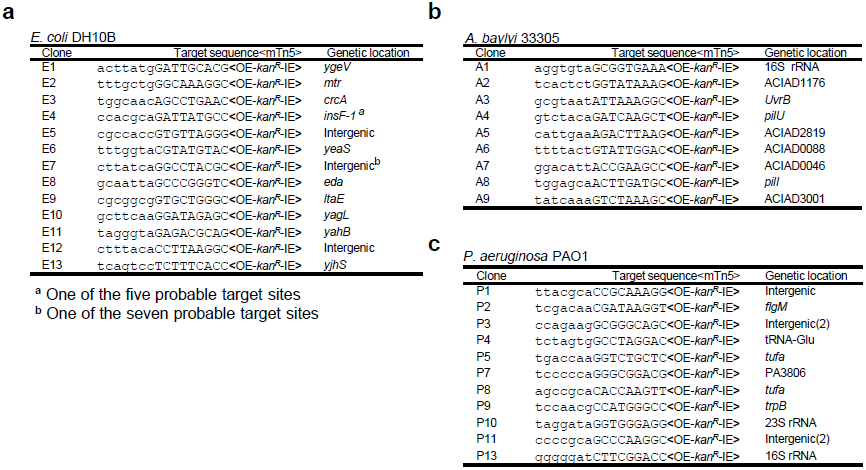
mTn5 insertion sites in different bacteria. (**a**) mTn5 insertion sites in *E. coli* DH10B. (**b**) mTn5 insertion sites in *A. baylyi* 33305. (**c**) mTn5 insertion sites in *P. aeruginosa* PAO1. Only the chromosomal sequences next to the OE are shown. The 9 bp duplicated sequences are shown in capital letters. Identical clones are shown only once, with numbers indicated in parenthesis. Either gene names or locus tags are given as genetic locations.

### Efficient mutagenesis of *Acinetobacter baylyi* and *P. aeruginosa* by a highly-active mTn5 transposon

Construct pSNC-mTn5 was then tested in two Gram-negative, capsule-bearing bacteria, *A. baylyi* 33305 and *P. aeruginosa* PAO1 [23, 24]. Comparing to *E. coli* DH10B, transformed *A. baylyi* 33305 and *P. aeruginosa* PAO1 appeared to lose the plasmid more easily in the absence of IPTG, with ~56.3% of *A. baylyi* and ~59.6% of *P. aeruginosa* retaining the plasmid after 24 h culture in LB media without antibiotics (Fig. 2e, f). Following IPTG induction, ~14.7% of *A. baylyi* and ~3.2% of *P. aeruginosa* retained the plasmid, suggesting that IPTG induced additional plasmid loss from these organisms, although their efficiencies were lower than that in DH10B cells. With sucrose counter selection, ~1.6 x 10^−7^ of *A. baylyi* remained Cam^R^, indicating that they contained the plasmid (Fig. 2e). Similarly, ~5.4 x 10^−7^ of *P. aeruginosa* cells were found to be Suc^R^Cam^R^ (Fig. 2f). These results suggest that IPTG and sucrose both contributed in reducing plasmid-bearing cells. In comparison, the percentages of Suc^R^Kan^R^ cells were 8.4 x 10^−6^ for *A. baylyi* and 3.4 x 10^−6^ for *P. aeruginosa*, suggesting that Tn5 transposition occurred in both organisms prior to plasmid loss. Colony restreaking showed that 100/100 Suc^R^Kan^R^ colonies are Suc^R^Cam^S^, suggesting that efficient plasmid loss had occurred following Tn5 transposition (~100% for both; Additional file 1: Figure S2a, c). Loss of plasmids was further confirmed by PCR tests (Additional file 1: Figure S2b, d). As observed in DH10B cells, Tn5 insertion also seemed to be random, as 9/9 *A. baylyi* and 13/15 *P. aeruginosa* mutants had different Tn5 insertion sites (Fig. 3b, c). The detection of identical mutants suggests that cell growth ensued following transposon transposition (Fig. 3c), which is common in different transposon mutagenesis assays [5, 6, 25].

### Construction of a Tn5 insertion library of *P. aeruginosa* using the highly-active mTn5 transposon

To determine whether we can construct a transposon insertion library of *P. aeruginosa* PAO1 with high diversity, ten pSNC-mTn5 transformants of the bacterium were cultured independently and then combined and induced with IPTG to initiate transposon mutagenesis. Following 24 h culture in LB media containing IPTG, ~6.4% of cells retained the plasmid (Additional file 1: Figure S3a). The frequencies of Suc^R^Cam^R^ and Suc^R^Kan^R^ cells in the IPTG-induced culture were found to be ~6.5 x 10^−7^ and ~3.5 x 10^−6^, respectively. Based on the total number of cells cultured and the frequency of Suc^R^Kan^R^Cam^S^ cells, the total diversity of the mTn5 insertion library was estimated to be ~1.3 x 10^7^, which covers the entire gene repertoire (5697) of *P. aeruginosa* PAO1 by ~2,238 times [24]. To our knowledge, the diversity of this transposon insertion library is bigger than the best transposon insertion library of PAO1 and related strains previously reported (Table 1) [5, 6, 9, 26-32]. Colony restreaking and PCR tests confirmed plasmid loss in the mutants (Additional file 1: Figure S3b, c), and 28/37 clones analyzed had different Tn5 insertion sites (Additional file 1: Figure S3d). Based on the percentage of independent clones in the library, its diversity is re-estimated to be ~1.0 x 10^7^.

**Table 1.**
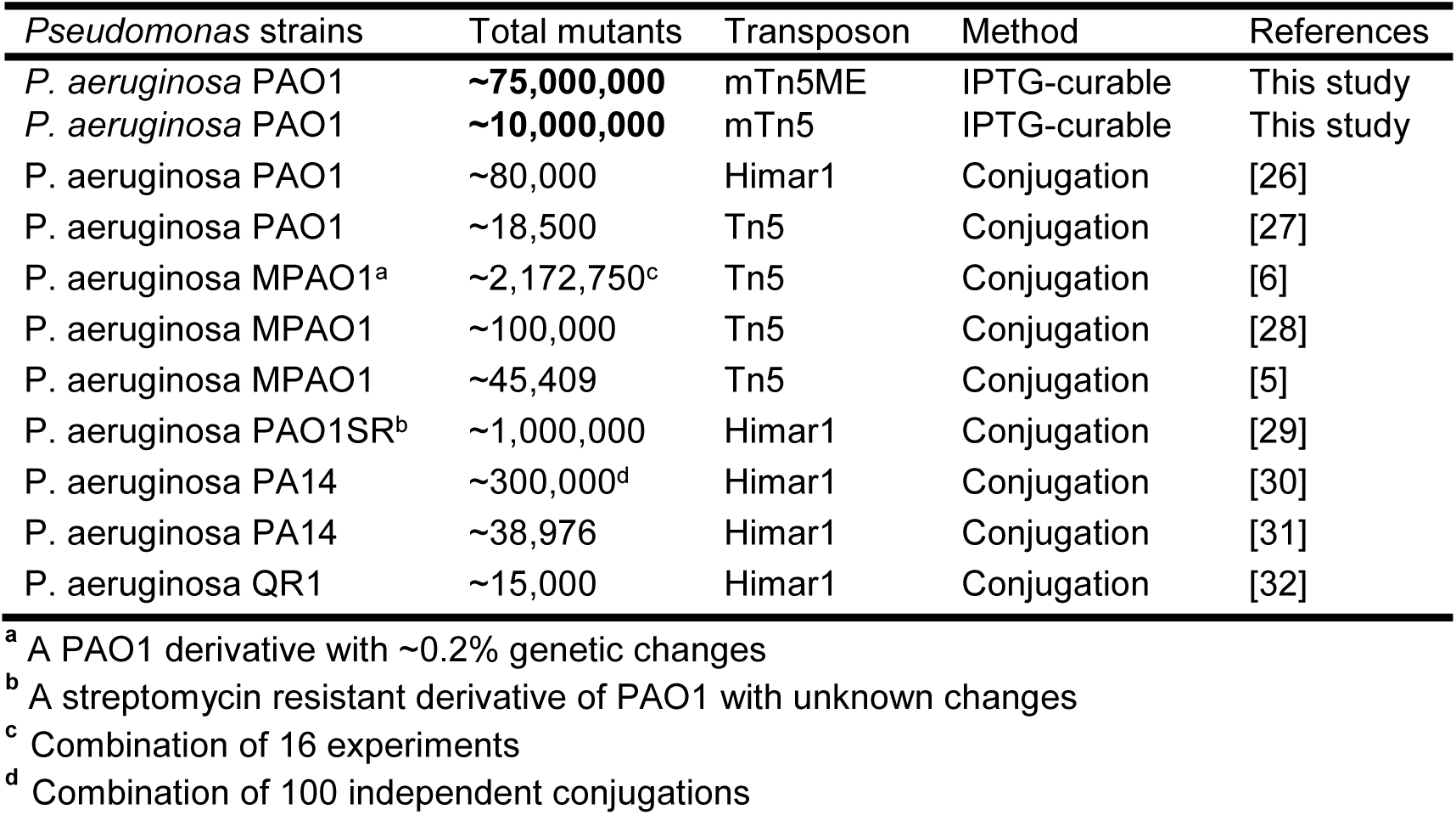
Comparison of transposon insertion libraries of *P. aeruginosa* strains

### An mTn5 with MEs enables generation of a *P. aeruginosa* mutant library with even higher diversity

To determine whether the efficiency of mTn5 transposition can be further improved, we replaced both OE and IE of the mTn5 with MEs (Fig. 4a). The new plasmid, pSNC-mTn5ME, was transformed into DH10B cells. Cell growth (or colony sizes) appeared to be normal, suggesting that basal-level transposition, if any, did not lead to obvious cellular toxicity, which was our initial concern. The behavior of the plasmid and Tn5 transposition efficiency were determined under the same conditions described above. Without IPTG induction, the plasmid remained relatively stable, as ~100% of the cells retained the plasmid (Cam^R^). After IPTG induction for 24 h, 1.1 x 10^−3^ of the cells retained the plasmid, suggesting that RepF overexpression caused efficient plasmid loss. With sucrose counter selection, ~2.4 x 10^−5^ cells remained Suc^R^Cam^R^. Interestingly, the frequency of Suc^R^Kan^R^ cells was found to be very high (~28.0%), indicating that mTn5ME is much more active than the non-ME version (~1200 folds). Restreaking of Suc^R^Kan^R^ colonies showed that they were all Kan^R^Cam^S^ (100/100) (Additional file 1: Figure S4a), and colony PCRs confirmed plasmid loss (10/10) (Additional file 1: Figure S4b). Sequence analysis of the transposon insertion junctions showed that all 13 Suc^R^Kan^R^ colonies analyzed had different Tn5 integration sites (Additional file 1: Figure S4c).

**Fig. 4.**
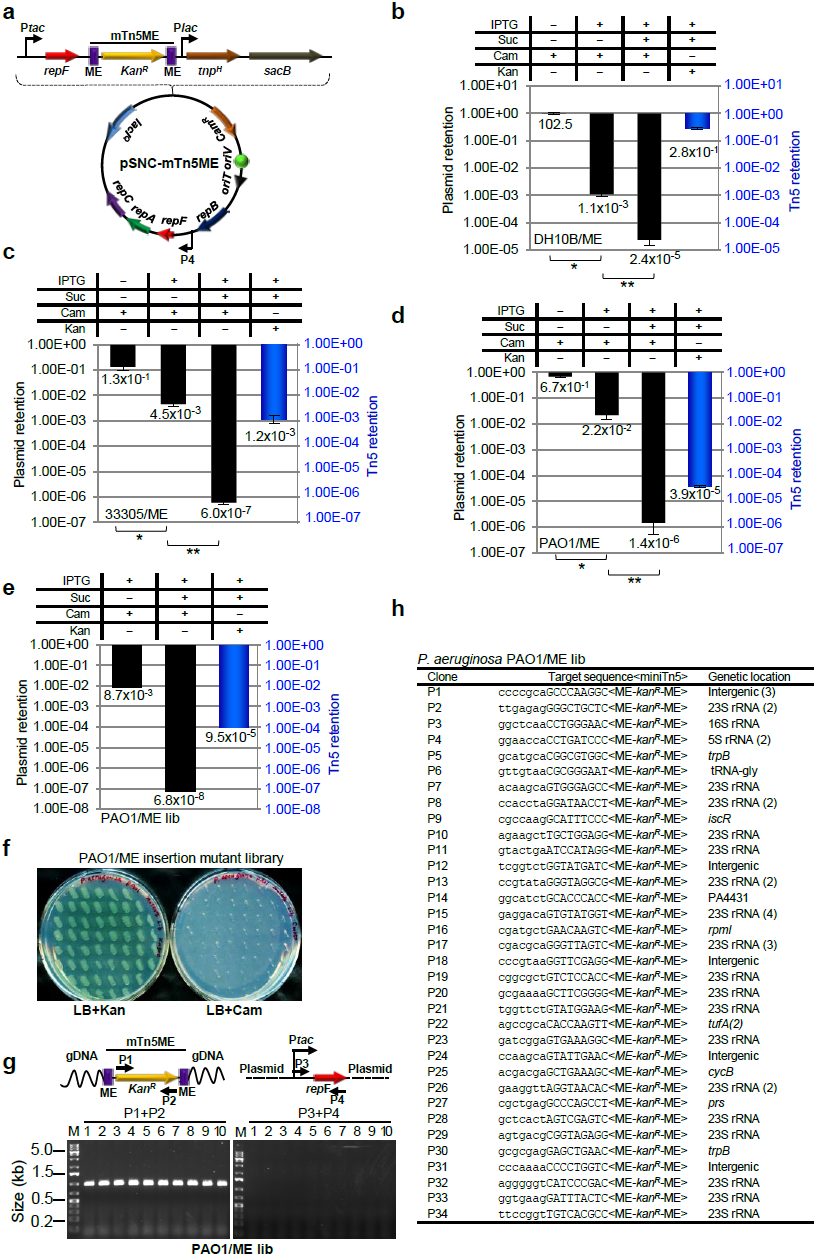
Generation of a *P. aeruginosa* insertion library with pSNC-mTn5ME. (**a**) Diagram of pSNC-mTn5ME, a derivative of pSNC-mTn5 that has MEs instead of OE and IE at the termini of mTn5. (**b**) Plasmid and transposon retention frequencies in *E. coli* DH10B. Results were average of three independent experiments, and bars represent mean ± SD (^*^p < 0.0001 and ^**^p = 0.0004 by unpaired t-test). Colony restreaking and PCR assays are shown in Additional file 1: Figure S4. (**c**) Plasmid and transposon retention frequencies in *A. baylyi*. Results were average of three independent experiments, and bars represent mean ± SD (^*^p = 0.0029 and ^**^p = 0.0006 by unpaired t-test). Colony restreaking and PCR assays are shown in Additional file 1: Figure S5. (**d**) Plasmid and transposon retention frequencies in *P. aeruginosa* PAO1. Results were average of three independent experiments, and bars represent mean ± SD (^*^p < 0.0001 and ^**^p = 0.0065 by unpaired t-test). Colony restreaking and PCR assays are shown in Additional file 1: Figure S6. (**e**) Plasmid and transposon retention frequencies in the *P. aeruginosa* PAO1 mutant library generated with pSNC-mTn5ME. (**f**) Colony restreaking. 100/100 Suc^R^Kan^R^ colonies of the mTn5ME library of *P. aeruginosa* were found to be Kan^R^Cam^S^. 50 are shown here. (**g**) Colony PCR of ten restreaked clones in (**f**) with the indicated primers. All were mTn5-positive and plasmid-negative. (**h**) Transposon insertion sites of 46 mutant clones from the mTn5ME insertion library of *P. aeruginosa*. Identical clones are shown only once, with their duplication numbers indicated in parenthesis.

We then determined whether construct pSNC-mTn5ME would also be more active in *A. baylyi* 33305 and in *P. aeruginosa* PAO1. For *A. baylyi*, the efficiencies of plasmid loss were higher for pSNC-mTn5ME than for pSNC-mTn5, both in the absence and presence of IPTG induction (Fig. 2e and 4c). Sucrose counter selection was highly effective for both constructs (Fig. 2e and 4c). As in *E. coli*, mTn5ME was found to be much more active than the non-ME version (~140 fold higher; Suc^R^Kan^R^ cells: ~1.2 x 10^−3^ for mTn5ME vs. ~8.4 x 10^−6^ for mTn5). Similarly, colony restreaking of Suc^R^Kan^R^ cells showed that 100/100 colonies are Kan^R^Cam^S^ (Additional file 1: Figure S5a), and plasmid loss was further verified by PCR (Additional file 1: Figure S5b). 9/14 colonies were found to have different Tn5 insertion sites (Additional file 1: Figure S5c). For PAO1, efficiencies of plasmid loss (±IPTG) were found to be similar for both pSNC-mTn5 and pSNC-mTn5ME (Fig. 2f and 4d), and sucrose counter selection was also effective for pSNC-mTn5ME (Fig. 4d). As in *E. coli* and in *A. baylyi*, mTn5ME was found to be more active than mTn5 in PAO1 (~11 fold higher; Suc^R^Kan^R^ cells: ~3.9 x 10^−5^ for mTn5ME vs. ~3.4 x 10^−6^ for mTn5) (Fig. 2f and 4d). In the colony restreaking assay, 100/100 Suc^R^Kan^R^ colonies were found to be Kan^R^Cam^S^ (Additional file 1: Figure S6a), and PCR assays further confirmed plasmid loss (10/10) (Additional file 1: Figure S6b). Sequence determination showed that 7/10 colonies tested had different Tn5 insertion sites (Additional file 1: Figure S6c).

We then determined whether pSNC-mTn5ME would be a better construct than pSNC-mTn5 for transposon saturation mutagenesis in *P. aeruginosa*. Ten transformants were randomly picked for Tn5 insertion library construction using the protocol described above. About 0.87% of cells retained the plasmid after IPTG induction, and the frequency of Suc^R^Cam^R^ cells was found to be 6.8 x 10^−8^. In comparison, the frequency of Suc^R^Kan^R^ cells was found to be ~9.5 x 10^−5^, suggesting that efficient mTn5ME transposition has occurred. Based on the total amount of cells cultured and the mTn5ME transposition efficiency (~9.5 x 10^−5^), the diversity of the mTn5ME insertion library was estimated to be 1.02 x 10^8^, which is ~3 logs larger than the best PAO1 transposon insertion library previously reported and ~2 logs larger than a Tn5 insertion library of *P. aeruginosa* MPAO1, a derivative of PAO1 with ~0.2% genetic variation [6, 33]. This new library is by far the biggest transposon insertion library of PAO1 and related species ever reported (Table 1). The size of our new library is enough to cover the entire gene repertoire of PAO1 by ~18,000 times. Colony restreaking (100) and PCR tests (10) confirmed plasmid loss in all the Suc^R^Kan^R^ clones analyzed (Fig. 4f, g), and 34/46 clones tested had different Tn5 insertion sites (Fig. 4h). Thus, the independent clones in the library (library diversity) are estimated to be ~7.5 x 10^7^.

## Discussion

We have developed a new transposon mutagenesis system that is efficient, regulatable, easy-to-use, and broadly useful. We believe it will be especially useful for functional genomics studies of Gram-negative bacteria that are difficult to transform and conjugate, such as certain capsule-containing bacteria, obligate anaerobes, and possibly obligate intracellular pathogens as well. The advantage of this method relies on the following features: (i) A broadly-functional plasmid replicon; (ii) Replication of the plasmid is regulated by IPTG; (iii) The inclusion of the *sacB* gene for counter selection; (iv) A highly-active/hyperactive transposon; (v) Regulatable expression of the hyperactive transposase gene; (vi) mTn5 and mTn5ME transposons insert almost completely randomly in different bacteria (Additional file 1: Figure S7) [34, 35]. In addition, the relatively small sizes of the RSF1010-based plasmids also facilitate their transformation and conjugation. Similar to transposon mutagenesis using *ts* plasmids, our system does not depend on efficient plasmid transformation and conjugation, and requires as few as one transformant or conjugated cell for transposon saturation mutagenesis. Using this new tool, we have generated a Tn5 transposon insertion library of *P. aeruginosa* PAO1 with a diversity of ~10^8^, which is ~2 logs larger than the best transposon insertion library of PAO1 and related *Pseudomonas* strains ever generated (Table 1). *P. aeruginosa* is an important opportunistic pathogen that frequently causes nosocomial infections and many of the strains are multidrug-resistant. The mutant PAO1 library we generated should also be valuable for *P. aeruginosa* pathogenesis studies.

To our knowledge, our plasmids are the only non-*ts*, conditional suicide plasmids used for transposon mutagenesis, and they replicate in a wide range of bacterial species [19]. In contrast, many *ts* mutant plasmids seem to have limited host ranges, either due to the limited host ranges of the parental plasmids, or due to the species-specificity of their *ts* phenotypes [10, 11, 14-17, 36]. In addition, *ts* plasmids often require prolonged incubation at high temperatures for plasmid curing, which can be harsh conditions for bacterial growth and survival, thus may lead to accumulation of adaptive genetic changes. Table S1 in Additional file 1 is a detailed comparison of our systems (pSNC-mTn5 and pSNC-mTn5ME) with various *ts* plasmid-based platforms that have been used for transposon mutagenesis in Gram-negative bacterial species, which clearly shows that our systems will be more broadly useful. In addition to their utilities in transposon mutagenesis, the IPTG-controlled conditional suicide plasmids that we developed should have many other applications, such as for allelic exchange or as curable vectors for delivering gene targeting systems, *e.g.*, TargeTrons, λ Red, RecET, *etc* [37, 38].

## Conclusion

In this work, we have developed a number of IPTG-controlled conditional suicide plasmids that contain the broad-host-range RSF1010 origin. Using one of the constructs to deliver a hyperactive mTn5 transposon, we showed that this system can be used for efficient mutagenesis of different bacterial species. As the assay condition is mild and the host range of the RSF1010 plasmid is extremely wide, we believe that our methodology will have broad applications in microbiology research.

## Methods

### Bacterial strains and growth conditions

*E. coli* DH10B was purchased from Invitrogen. *A. baylyi* (ATCC 33305) and *P. aeruginosa* PAO1 (ATCC BAA-47) were purchased from ATCC. Unless stated otherwise, all the strains were grown at 37°C in Luria Broth (LB) liquid media with agitation at 200 rpm or on LB plates with 1.5% agar. For *sacB* counter selection, we used LBNS plates (LB no salt: 1% Tryptone, 0.5% yeast extract and 1.5% agar) supplemented with 10% sucrose (Fisher Scientific). Appropriate antibiotics and concentrations were used to select for bacterial cells that are antibiotic resistant. *E. coli* DH10B: chloramphenicol (Cam; Gold Biotechnology), 25 µg/ml; kanamycin (Kan; Fisher Scientific), 50 µg/ml. *A. baylyi*: Cam, 10 µg/ml; Kan, 10 µg/ml. *P. aeruginosa* PAO1: Cam, 250 µg/ml; Kan, 500 µg/ml.

### Plasmid construction

To construct plamsid pMMB-*repF*, we PCR-amplified the *repF* gene from pMMB208 [18]. The PCR fragment was digested with *Hin*dIII and *Pst*I, and inserted at the corresponding sites of pMMB208, downstream of the *tac* promoter (P*tac*). To construct plasmid pMMB-*repF*/*sacB*, the *sacB* gene and its promoter were PCR amplified from plasmid pRE112 [22] and inserted between the unique *Sac*I and *Kpn*I sites of pMMB-*repF*.

To construct plasmids pMMB-*repA*K42A and pMMB-*repA*D139A, *repA* genes containing K42A and D139A mutations were generated in two-step PCRs from plasmid pMMB208 [21]. The mutant genes were cloned between the *Hin*dIII and *Pst*I sites of pMMB208.

Plasmid pSNC-mTn5 was constructed in multiple steps. First, plasmid pUT-mTn5Km/lacEZ was constructed from plasmid pUT-mTn5Km [4]. It contains a *lac* promoter-driven hyperactive transposase gene (*tnp*^*H*^) that has E54K, M56A and L372P mutations [3]. In addition, inside the mTn5 transposon, the inverted repeats flanking the kanamycin resistance marker (*Kan*^*R*^) were deleted [4]. The entire mTn5 cassette of pUT-mTn5Km/lacEZ, which contains the *Kan*^*R*^-mTn5 transposon and P*lac*-*tnp*^*H*^, was then PCR amplified and cloned at the *Xba*I site of pMMB-*repF*/*sacB*, resulting in plasmid pSNC-mTn5. It has an OE and an IE at the termini of the mTn5. Plasmid pSNC-mTn5ME was derived from pSNC-mTn5 by replacing both OE and IE with MEs.

### Characterization of IPTG-induced plasmid loss and Transposon mutagenesis

To test IPTG-induced plasmid loss of pMMB-*repF*, pMMB-*repA*K42A and pMMB-*repA*D139A, single colonies of *E. coli* DH10B cells transformed with the plasmids were inoculated into 5 ml LB+Cam media and cultured at 37°C for ~14 hours (h). After measuring OD600, 1 ml of each culture was pelleted by centrifugation and washed with 500 µl of fresh LB to remove antibiotics. Cells were then resuspended in 1 ml LB. An aliquot was added to 5 ml LB (final OD600 = 0.001) with and without 1 mM IPTG and cultured at 37°C for 24 h. 1 ml of the IPTG-induced samples was then pelleted, washed with 500 µl LB, and resuspended in 1 ml LB. Serial dilutions of the samples (±IPTG) were plated on LB and LB+Cam plates to evaluate plasmid loss. Plasmid retention frequencies were calculated as ratios of cfu (colony forming units) on LB+Cam plates and those on LB plates.

To perform transposon mutagenesis in *E. coli* DH10B, single colonies of pSNC-mTn5 and pSNC-mTn5ME transformants were cultured in 5 ml LB+Cam+Kan media overnight at 37°C. Cells were then pelleted and washed as above to remove antibiotics, and an aliquot was inoculated to 5 ml LB (final OD600 = 0.001) in a 14 ml culture tube and grown at 37°C for 24 h with and without 1 mM IPTG induction. A 1 ml aliquot of the IPTG-induced samples was then pelleted, washed with 500 µl LBNS, and resuspended in 1 ml LBNS. Serial dilutions of the samples (±IPTG) were plated on LB and LB+Cam plates to evaluate plasmid loss. The IPTG induced samples were also plated on LBNS+10% sucrose and LBNS+10% sucrose+Cam plates to estimate percentage of plasmid-retaining cells in the presence of sucrose counter selection; and LBNS+10% sucrose+Kan plates to select for transposition events. Plasmid retention frequencies (PRF) were calculated as the following: (1) −IPTG: (cfu on LB+Cam)/(cfu on LB); (2) +IPTG: (cfu on LB+Cam)/(cfu on LB); (3) +IPTG+Suc: (cfu on LBNS+Suc+Cam)/(cfu on LBNS+Suc). Transposon retention frequencies (TRF) were calculated as the following: +IPTG+Suc: (cfu on LBNS+Suc+Kan)/(cfu on LBNS+Suc). mTn5 (or mTn5ME) transposition frequencies were calculated as TRF+IPTG+Suc – PRF+IPTG+Suc, which essentially equals to TRF+IPTG+Suc if the background (PRF+IPTG+Suc) is low. The same protocol, except for the concentrations of antibiotics (indicated above) and IPTG (10 mM for PAO1), was followed to perform transposon mutagenesis in *P. aeruginosa* PAO1.

Similarly, to perform transposon mutagenesis in *A. baylyi* 33305, single colonies of pSNC-mTn5 and pSNC-mTn5ME transformants were cultured in 5 ml LB+Cam+Kan media overnight at 37°C. Cells were pelleted and washed as for *E. coli* and *P. aeruginosa*. Then, an aliquot was inoculated to 100 ml LB (final OD600 = 0.001) in a baffled flask. The cultures were shaken vigorously (~250 rpm) at 37°C for 24 h with and without 10 mM IPTG induction. A 1 ml aliquot of the IPTG-induced samples was then pelleted, washed with 500 µl LBNS, and resuspended in 1 ml LBNS. Serial dilutions of the samples (±IPTG) were then plated on appropriate plates to evaluate plasmid loss and mTn5 (or mTn5ME) transposition as in the assays for *E. coli* and for *P. aeruginosa*.

To verify plasmid loss in cells with potential transposition events, 100-150 Suc^R^Kan^R^ colonies in each assay were then restreaked on LB+Kan and LB+Cam plates. In addition, presence of the transposon and the plasmid was determined by colony PCRs in a 25 µl reaction containing 25 mM TAPS-HCl (pH 9.3), 50 mM KCl, 2 mM MgCl2, 1 mM β-mercaptoethanol, 1x GC enhancer, 0.2 mM dNTPs, 0.1 µl of Q5 polymerase (2 u/µl; NEB), 1 µl of resuspended cells, and 150 ng each of the primers (final concentration = ~0.5 µM; see figure legends and Table S2 in Additional file 1 for oligos used). PCRs were performed using the following condition: 1x (94°C, 2 minutes); 25x (94°C, 30 seconds; 50°C, 30 seconds; 72°C, 1 minute); 1x (72°C, 10 minutes); 1x (4°C, hold).

### Determination of transposon insertion sites

Transposon insertion sites in bacterial chromosomes were determined by arbitrarily primed PCR, in which transposon junctions were amplified in two steps [5, 39]. Bacterial cells were resuspended in 10-20 µl of deionized water and 1 µl was used directly as the PCR template. In the first PCR step, the reaction was performed using a specific primer annealing to the transposon region (Tn5Km1) and a semi-degenerate primer (BDC1) that anneals to many sites on the bacterial chromosome. In the second step, aliquots of the first-round PCR products were amplified using a primer annealing to the transposon region (Tn5Km2), slightly closer to the insertion junction, and a non-degenerate primer (BDC2) that anneals to the constant region of the BDC1-derived sequence. PCRs were carried out under the conditions described above. PCR products from Step 2 were resolved in a 2% agarose gel and major products were gel-purified for sequencing to determine Tn5 insertion sites.

### Construction of transposon insertion libraries of *P. aeruginosa* PAO1

To construct an mTn5 (or mTn5ME) insertion library of *P. aeruginosa* PAO1, plasmid pSNC-mTn5 (or pSNC-mTn5ME) was first electroporated into the bacterial cells. Ten transformants were cultured independently in 5 ml LB+Cam+Kan media at 37°C for ~14 hours. Equal amount of each sample (equivalent to 0.5 OD600 x 1 ml) was then combined, pelleted, washed with 500 µl LB, and the pellet was resuspended in 1 ml LB. An aliquot of the mixture was then inoculated into 500 ml LB supplemented with 10 mM IPTG in a baffled flask (final OD600 = 0.01) and shaken vigorously (300 rpm) at 37°C for 24 h to perform transposon mutagenesis. The cells were then pelleted by centrifugation and washed with 250 ml LBNS medium. The pellet was resuspended in 50 ml LBNS medium and serial dilutions were plated on LB, LB+Cam, LBNS+10% sucrose, LBNS+10% sucrose+Cam, and LBNS+10% sucrose+Kan plates to determine plasmid loss, mTn5 (mTn5ME) transposition, and total library diversity. Arbitrary PCR and DNA sequencing were then performed to determine Tn5 insertion sites.

### Determination of mTn5 and mTn5ME target site preferences in *P. aeruginosa* PAO1

To determine if mTn5 and mTn5ME have any target site preferences in *P. aeruginosa* PAO1, we generated sequence logos of their insertion sites in the bacterium using the WebLogo server (https://weblogo.berkeley.edu/logo.cgi). In total, 40 mTn5 insertion sites and 41 mTn5ME insertion sites were used for the analysis.

## Abbreviations

IPTG: Isopropyl β-D-1-thiogalactopyranoside;
*ts*: Temperature-Sensitive;
bp: base pair;
OE: Outside End;
IE: Inside End;
ME: Mosaic End;
OD: Optical Density;
LB: Luria Broth;
LBNS: LB no salt;
Cam: Chloramphenicol;
Kan: kanamycin.

## Acknowledgements

We thank Drs. Mark McIntosh, David Pintel and Donald H. Burke for helpful discussions.

## Availability of data and materials

The dataset supporting the conclusion of this article are available from the corresponding author on reasonable request.

## Authors’ contributions

SSN, HG conceived the study and designed the experiments; SSN, JH, SYZ, JZ, AS, LBG, CVS, FR, performed the experiments; SSN and HG, wrote the manuscript. All authors have read and approved the manuscript.

## Ethics approval and consent to participate

Not applicable.

## Consent for publication

Not applicable.

## Additional files

**Additional file 1: Figure S1.** Tn5 transposons. (**a**) Full-length Tn5. The full-length Tn5 contains two inverted IS50 elements at its ends. Only one of them encodes an active Tnp and an Inh (Inhibitor of Tnp). *Kan*^*R*^, kanamycin-resistance gene; *Str*^*R*^, streptomycin-resistance gene; and *Ble*^*R*^, bleomycin-resistance gene. (**b**) mTn5s. Top, an mTn5 with an OE and an IE at the termini. Bottom, an mTn5 with MEs at the ends. (**c**) Comparison of OE, IE and ME, with their polymorphisms highlighted in red.

**Additional file 1: Figure S2.** Confirmation of mTn5 transposition events in *A. baylyi* and in *P. aeruginosa*. (**a**) Colony restreaking assay of *A. baylyi*. 100 Suc^R^Kan^R^ colonies of *A. baylyi* were restreaked on LB+Kan and LB+Cam plates, and all were found to be Kan^R^Cam^S^. 50 are shown here. (**b**) Colony PCR of 10 restreaked *A. baylyi* clones with the indicated primers. All were mTn5-positive and plasmid-negative. (**c**) Colony restreaking assay of *P. aeruginosa*. 100 Suc^R^Kan^R^ colonies of *P. aeruginosa* were restreaked on LB+Kan and LB+Cam plates, and all were found to be Kan^R^Cam^S^. 50 are shown here. (**d**) Colony PCR of ten restreaked *P. aeruginosa* clones with primers indicated in the diagram. All were mTn5-positive and plasmid-negative.

**Additional file 1: Figure S3.** A transposon insertion library of *P. aeruginosa* PAO1 generated with pSNC-mTn5. (**a**) Plasmid and transposon retention frequencies of the mTn5 insertion library of *P. aeruginosa* PAO1. (**b**) Colony restreaking assay. 100 random Suc^R^Kan^R^ colonies were restreaked on LB+Kan and LB+Cam plates. 100/100 were found to be Kan^R^Cam^S^ and 50 are shown here. (**c**) Colony PCR of ten restreaked clones in (**b**). All were found to be mTn5-positive and plasmid-negative. (**d**) mTn5 insertion sites of 37 mutant clones from the transposon insertion library of *P. aeruginosa*. Identical clones are only shown once, and their numbers are indicated in parenthesis.

**Additional file 1: Figure S4.** Confirmation of mTn5ME transposition events in *E. coli* DH10B. (**a**) Colony restreaking. 100 random Suc^R^Kan^R^ colonies of *E. coli* were restreaked on LB+Kan and LB+Cam plates. 100/100 were found to be Kan^R^Cam^S^ and 50 restreaked colonies are shown here. (**b**) Colony PCR of ten restreaked clones in (**a**). All were found to be Tn5-positive and plasmid-negative. (**c**) Tn5 insertion sites of 13 independent DH10B clones.

**Additional file 1: Figure S5.** Confirmation of mTn5ME transposition events in *A. baylyi* 33305. (**a**) Colony restreaking. 100 random Suc^R^Kan^R^ colonies of *A. baylyi* were restreaked on LB+Kan and LB+Cam plates. 100/100 were found to be Kan^R^Cam^S^ and 50 restreaked colonies are shown here. (**b**) Colony PCR of ten restreaked clones in (**a**). All were found to be Tn5-positive and plasmid-negative. (**c**) Sequence analysis shows that 9/14 *A. baylyi* clones had different Tn5 insertion sites.

**Additional file 1: Figure S6.** Confirmation of mTn5ME transposition events in *P. aeruginosa* PAO1. (**a**) Colony restreaking. 100 random Suc^R^Kan^R^ colonies of *P. aeruginosa* PAO1 were restreaked on LB+Kan and LB+Cam plates. 100/100 were found to be Kan^R^Cam^S^ and 50 restreaked colonies are shown here. (**b**) Colony PCR of ten restreaked clones in (**a**). All were found to be Tn5-positive and plasmid-negative. (**c**) Sequence analysis shows that 7/10 *P. aeruginosa* clones had different Tn5 insertion sites.

**Additional file 1: Figure S7.** Target site preferences of mTn5 and mTn5ME in *P. aeruginosa*. (**a**) Sequence logo of mTn5 insertion sites generated with WebLogo. 40 target sequences were analyzed. The 9 bp duplicated sequences adjacent to the OE are shown. There is a slight preference for certain nucleotides at several positions. (**b**) Sequence logo of mTn5ME insertion sites. 41 target sequences were analyzed. The 9 bp duplicated sequences adjacent to an ME are shown. It appears that mTn5ME has less nucleotide preference at the duplicated target sequence than mTn5 in *Pseudomonas*.

**Additional file 1: Table S1.** Comparison of conditional suicide vector-based transposon mutagenesis strategies used in Gram-negative bacteria.

**Additional file 1: Table S2.** List of primers used.

## References

1. Phadnis SH, Berg DE: Identification of base pairs in the outside end of insertion sequence IS50 that are needed for IS50 and Tn5 transposition. Proc Natl Acad Sci U S A. 1987;84(24):9118–9122.

2. Reznikoff WS: Transposon Tn5. Annu Rev Genet. 2008;42:269–286.

3. Goryshin IY, Reznikoff WS: Tn5 in vitro transposition. J Biol Chem. 1998;273(13):7367–7374.

4. de Lorenzo V, Herrero M, Jakubzik U, Timmis KN: Mini-Tn5 transposon derivatives for insertion mutagenesis, promoter probing, and chromosomal insertion of cloned DNA in gram-negative eubacteria. J Bacteriol. 1990;172(11):6568–6572.

5. Jacobs MA, Alwood A, Thaipisuttikul I, Spencer D, Haugen E, Ernst S, Will O, Kaul R, Raymond C, Levy R et al: Comprehensive transposon mutant library of Pseudomonas aeruginosa. Proc Natl Acad Sci U S A. 2003;100(24):14339–14344.

6. Lee SA, Gallagher LA, Thongdee M, Staudinger BJ, Lippman S, Singh PK, Manoil C: General and condition-specific essential functions of Pseudomonas aeruginosa. Proc Natl Acad Sci U S A. 2015;112(16):5189–5194.

7. Metzger M, Bellemann P, Schwartz T, Geider K: Site-directed and transposon-mediated mutagenesis with pfd-plasmids by electroporation of Erwinia amylovora and Escherichia coli cells. Nucleic Acids Res. 1992;20(9):2265–2270.

8. Leahy JG, Jonesmeehan JM, Colwell RR: Transformation of Acinetobacter-Calcoaceticus Rag-1 by Electroporation. Can J Microbiol. 1994;40(3):233–236.

9. Shan Z, Xu H, Shi X, Yu Y, Yao H, Zhang X, Bai Y, Gao C, Saris PE, Qiao M: Identification of two new genes involved in twitching motility in Pseudomonas aeruginosa. Microbiology. 2004;150(Pt 8):2653–2661.

10. Sasakawa C, Yoshikawa M: A series of Tn5 variants with various drug-resistance markers and suicide vector for transposon mutagenesis. Gene. 1987;56(2-3):283–288.

11. Harayama S, Tsuda M, Iino T: Tn1 insertion mutagenesis in Escherichia coli K-12 using a temperature-sensitive mutant of plasmid RP4. Mol Gen Genet. 1981;184(1):52–55.

12. Le Breton Y, Mohapatra NP, Haldenwang WG: In vivo random mutagenesis of Bacillus subtilis by use of TnYLB-1, a mariner-based transposon. Appl Environ Microbiol. 2006;72(1):327–333.

13. Stubbendieck RM, Straight PD: Linearmycins Activate a Two-Component Signaling System Involved in Bacterial Competition and Biofilm Morphology. J Bacteriol. 2017;199(18).

14. Maier TM, Pechous R, Casey M, Zahrt TC, Frank DW: In vivo Himar1-based transposon mutagenesis of Francisella tularensis. Appl Environ Microbiol. 2006;72(3):1878–1885.

15. Rholl DA, Trunck LA, Schweizer HP: In vivo Himar1 transposon mutagenesis of Burkholderia pseudomallei. Appl Environ Microbiol. 2008;74(24):7529–7535.

16. Pelicic V, Jackson M, Reyrat JM, Jacobs WR, Jr., Gicquel B, Guilhot C: Efficient allelic exchange and transposon mutagenesis in Mycobacterium tuberculosis. Proc Natl Acad Sci U S A. 1997;94(20):10955–10960.

17. Choi KH, Mima T, Casart Y, Rholl D, Kumar A, Beacham IR, Schweizer HP: Genetic tools for select-agent-compliant manipulation of Burkholderia pseudomallei. Appl Environ Microbiol. 2008;74(4):1064–1075.

18. Morales VM, Backman A, Bagdasarian M: A series of wide-host-range low-copy-number vectors that allow direct screening for recombinants. Gene. 1991;97(1):39– 47.

19. Meyer R: Replication and conjugative mobilization of broad host-range IncQ plasmids. Plasmid. 2009;62(2):57–70.

20. Maeser S, Scholz P, Otto S, Scherzinger E: Gene F of plasmid RSF1010 codes for a low-molecular-weight repressor protein that autoregulates expression of the repAC operon. Nucleic Acids Res. 1990;18(21):6215–6222.

21. Ziegelin G, Niedenzu T, Lurz R, Saenger W, Lanka E: Hexameric RSF1010 helicase RepA: the structural and functional importance of single amino acid residues. Nucleic Acids Res. 2003;31(20):5917–5929.

22. Edwards RA, Keller LH, Schifferli DM: Improved allelic exchange vectors and their use to analyze 987P fimbria gene expression. Gene. 1998;207(2):149–157.

23. Barbe V, Vallenet D, Fonknechten N, Kreimeyer A, Oztas S, Labarre L, Cruveiller S, Robert C, Duprat S, Wincker P et al: Unique features revealed by the genome sequence of Acinetobacter sp. ADP1, a versatile and naturally transformation competent bacterium. Nucleic Acids Res. 2004;32(19):5766–5779.

24. Stover CK, Pham XQ, Erwin AL, Mizoguchi SD, Warrener P, Hickey MJ, Brinkman FS, Hufnagle WO, Kowalik DJ, Lagrou M et al: Complete genome sequence of Pseudomonas aeruginosa PAO1, an opportunistic pathogen. Nature. 2000;406(6799):959–964.

25. Wang N, Ozer EA, Mandel MJ, Hauser AR: Genome-wide identification of Acinetobacter baumannii genes necessary for persistence in the lung. MBio. 2014;5(3):e01163–01114.

26. Withers TR, Yin Y, Yu HD: Identification of novel genes associated with alginate production in Pseudomonas aeruginosa using mini-himar1 mariner transposon-mediated mutagenesis. J Vis Exp. 2014;85:51346.

27. Lewenza S, Falsafi RK, Winsor G, Gooderham WJ, McPhee JB, Brinkman FS, Hancock RE: Construction of a mini-Tn5-luxCDABE mutant library in Pseudomonas aeruginosa PAO1: a tool for identifying differentially regulated genes. Genome Res. 2005;15(4):583–589.

28. Gallagher LA, Shendure J, Manoil C: Genome-scale identification of resistance functions in Pseudomonas aeruginosa using Tn-seq. MBio. 2011;2(1):e00315–00310.

29. Wong SM, Mekalanos JJ: Genetic footprinting with mariner-based transposition in Pseudomonas aeruginosa. Proc Natl Acad Sci U S A. 2000;97(18):10191–10196.

30. Skurnik D, Roux D, Aschard H, Cattoir V, Yoder-Himes D, Lory S, Pier GB: A comprehensive analysis of in vitro and in vivo genetic fitness of Pseudomonas aeruginosa using high-throughput sequencing of transposon libraries. PLoS Pathog. 2013;9(9):e1003582.

31. Liberati NT, Urbach JM, Miyata S, Lee DG, Drenkard E, Wu G, Villanueva J, Wei T, Ausubel FM: An ordered, nonredundant library of Pseudomonas aeruginosa strain PA14 transposon insertion mutants. Proc Natl Acad Sci U S A. 2006;103(8):2833– 2838.

32. Seet Q, Zhang LH: Anti-activator QslA defines the quorum sensing threshold and response in Pseudomonas aeruginosa. Mol Microbiol. 2011;80(4):951–965.

33. Klockgether J, Munder A, Neugebauer J, Davenport CF, Stanke F, Larbig KD, Heeb S, Schock U, Pohl TM, Wiehlmann L et al: Genome diversity of Pseudomonas aeruginosa PAO1 laboratory strains. J Bacteriol. 2010;192(4):1113–1121.

34. Green B, Bouchier C, Fairhead C, Craig NL, Cormack BP: Insertion site preference of Mu, Tn5, and Tn7 transposons. Mob DNA. 2012;3(1):3.

35. Goryshin IY, Miller JA, Kil YV, Lanzov VA, Reznikoff WS: Tn5/IS50 target recognition. Proc Natl Acad Sci U S A. 1998;95(18):10716–10721.

36. Maier TM, Havig A, Casey M, Nano FE, Frank DW, Zahrt TC: Construction and characterization of a highly efficient Francisella shuttle plasmid. Appl Environ Microbiol. 2004;70(12):7511–7519.

37. Enyeart PJ, Mohr G, Ellington AD, Lambowitz AM: Biotechnological applications of mobile group II introns and their reverse transcriptases: gene targeting, RNA-seq, and non-coding RNA analysis. Mob DNA. 2014;5(1):2.

38. Court DL, Sawitzke JA, Thomason LC: Genetic engineering using homologous recombination. Annu Rev Genet. 2002;36:361–388.

39. Saavedra JT, Schwartzman JA, Gilmore MS: Mapping Transposon Insertions in Bacterial Genomes by Arbitrarily Primed PCR. Curr Protoc Mol Biol. 2017;118:15.15.11–15.15.15.

